# Dynamic Phase Synchrony based Ranked Spatio-Temporal Clustering for Tracking Time-Resolved Functional Brain Networks

**DOI:** 10.1101/172981

**Authors:** Priya Aggarwal, Anubha Gupta

## Abstract

Of late, there has been a growing interest in studying brain networks, particularly, for understanding spontaneous temporal changes in functional brain networks. Recently, phase synchrony based methods have been proposed to track instantaneous time-resolved functional connectivity without any need of windowing the data. This paper extends one such recently used phase synchrony measure in two steps. First, multiple temporal models are built from four-mode tensor that are further clustered to detect dynamic brain network communities. This clustering is based on spatio-temporal data and hence, is named as *Spatio-Temporal Clustering* (STC). Second, a method is proposed to rank all the communities allowing the proposed model to deal with multiple communities of differing time evolution. This helps in the comparison of network communities, especially, when available communities are too dense to provide relevant information for comparison. The ranking of communities allows for the dimensionality reduction of communities, while still maintaining the key brain networks. Intrinsic time-varying functional connectivity has been investigated for large scale brain networks, including default-mode network (DMN), visual network (VN), cognitive control network (CCN), auditory network (AN), etc. The proposed method provides a new complementary tool to investigate dynamic network states at a high temporal resolution and is tested on resting-state functional MRI data of 26 typically developing controls (TDC) and 35 autism spectrum disorder (ASD) subjects. Simulation results demonstrate that ASD subjects have altered dynamic brain networks compared to TDC.

## 1. Introduction

Functional connectivity analysis in brain is an important method in neuroscience for functional brain networks discovery Bullmore & Sporns (2009); Friston (2011); Biswal et al. (1995)). The functional Magnetic Resonance Imaging (fMRI) provides information about large-scale functional brain networks such as visual network (VN), somato-motor Network (SMN), auditory network (AN), cognitive control network (CCN), defaultmode network (DMN) etc. Fox & Raichle (2007); Doucet et al. (2011); Hacker et al. (2013); Power et al. (2011); Thomas Yeo et al. (2011). The conventional static connectivity analysis assume brain networks to be constant during the entire time duration of an fMRI scan session and therefore, fails to account for the time-varying nature of brain networks. Increasingly, dynamic functional connectivity (dFC) is being used to capture and characterize time-varying brain networks Chang & Glover (2010); Sakoğlu et al. (2010); Kiviniemi et al. (2011) as supported by a growing body of research suggesting for dynamic reconfiguration of brain networks Jones et al. (2012); Allen et al. (2014); Leonardi et al. (2013); Hutchison et al. (2013b); Zalesky et al. (2014); Calhoun et al. (2014).

A sliding-window (SW) approach is commonly used on fMRI time series data to elucidate dynamic aspects of functional brain networks (see Hutchison et al. (2013a) for review). In this approach, brain networks are assumed to be time-invariant within a predefined window duration. This window is subsequently shifted either by one or many time points to estimate temporal changes in brain networks. However, this method is suboptimal due to the strong dependence of observations in dFC on the window size and the overlap between windows Shakil et al. (2016); Hindriks et al. (2016). Moreover, there may be abrupt changes in the functional connectivity at different time points. Hence, fixed window assumption of SW analysis may not necessarily hold true Monti et al. (2017).

Besides considering conventional SW approach, other attempts have been made to find instantaneous patterns of dFC without windowing and hence, with no need of struggling with choosing the right window length Glerean et al. (2012); Ponce-Alvarez et al. (2015); Omidvarnia et al. (2016); Demirta et al. (2016); Córdova-Palomera et al. (2017). Phase difference between the time series of regions is one such instantaneous measure of dFC and is commonly known as phase synchrony. This approach computes connectivity at every time point and hence, alleviates the need of choosing window length. Moreover, this method allows extraction of dynamic functional connectivity at higher temporal resolution compared to the state-of-the-art sliding-window approach. To date, several studies have utilized phase synchrony measure to compute dFC in resting state fMRI Glerean et al. (2012); Ponce-Alvarez et al. (2015); Omidvarnia et al. (2016); Demirta et al. (2016); Córdova-Palomera et al. (2017).

In the present study, we seek whole-brain dynamic brain networks’ topology using instantaneous phase measure computed using the Hilbert transform Glerean et al. (2012). In this approach, first, a signal is bandpass filtered. Next, instantaneous phase is estimated from the signal’s analytic form obtained from the Hilbert transform. We propose to use this measure to build dynamic brain networks.

After obtaining dFC, next task is to identify communities of densely connected regions and to study their temporally dynamic profiles from multiple temporal snapshots of phase synchrony matrices. Related work of identifying communities using phase-synchrony based time-varying matrices is implemented by decomposing a three-mode tensor of regional connectivity at different times and then analyzing and linking the changes in the community structures Ponce-Alvarez et al. (2015). For example, in Ponce-Alvarez et al. (2015), tensor decomposition based method on three-mode tensor (region × region × time) is applied to obtain a few significant components. Each component consists of three vectors, namely, loading vectors. The first two loading vectors are related to regions and are used to generate communities. The third loading vector contains temporal information of a community’s time-varying dynamic profile. We observe that direct utilization of tensor decomposition fails to provide a good model for dynamic brain network communities due to random fluctuation of loading vectors related to regions. Interpretations of these vectors does not provide consistent results.

To overcome the above issue, we propose a Spatio-Temporal Clustering (STC) framework based on the temporal models learned from tensor decomposed loading vectors. Our framework identifies multiple clusters from each temporal model. This results in many clusters, with a time loading vector for each cluster. With a number of identified clusters, we rank them using our proposed Combined Cluster Score (CCS) based on edge values between regions in a cluster and corresponding temporal loading vector. Thus, we obtain multiple dynamic brain networks communities. We will formalize our proposed model in the next few sections.

Major contributions of this work are as follows:

- We provide a spatio-temporal clustering framework to detect dynamic brain network communities using K-means clustering and Silhouette criterion. Our model allows a region to belong to multiple network communities and hence, provides a novel method to detect overlapping dynamic brain network communities.
- We provide a method to rank all the clusters that allows our model to deal with a large number of clusters having different temporal evolution. This helps with the comparison of network communities, especially when available communities are too dense to provide relevant information for comparison.
- We differentiate subjects suffering with autism spectrum disorder from healthy controls using the proposed dynamic functional connectivity analysis. In addition, we provide a new analysis tool to explore dynamic functional connectivity and characterize relationship between regions of interests (ROIs) without explicitly defining window length as required in the conventional ‘sliding window’ based dynamic functional connectivity analysis.

The organization of this paper is as follows: Section 2 describes materials used in this paper. Section 3 explains the steps to learn and track dynamic brain network communities and their temporal evolution profile. Section 4 reports experiments with real data. In the end, discussion and conclusions are presented in section 5 and 6, respectively.

*Notations:* We use lowercase boldface letters for vectors (e.g. **a**), capital boldface letters for matrices (e.g. **A**), italics letters for scalars (e.g. *a* or A), and Euler script letters for tensors (e.g. 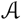).

## 2. Materials

### 2.0.1. Data Acquisition

In this paper, we have used publicly available Autism dataset contributed by Georgetown University, a collection site of Autism Brain Image Data Exchange II (ABIDEII) http://fcon_1000.projects.nitrc.org/indi/abide/. To exclude effects of multiple sites, we analyzed data collected at a single site (here Georgetown University). This site was choosen because of the availability of a larger number of adolescent males with ASD. The fMRI data of 55 healthy subjects (Typically Developing Controls (TDC)) (8.1’13.8 years) and 51 Autism Spectrum Disorder (ASD) subjects (8.1’13.9 years) is available. A total of 152 brain volume fMRI data has been collected with Echo Time (TE) equal to 30 milli-seconds (ms) and Repetition Time (TR) equal to 2000 ms. Each brain volume consists of an acquisition of 43 brain slices of size 64 × 64. A three-dimensional structural magnetization prepared rapid gradient echo *T*1-weighted image is acquired for every subject with TR = 2530 ms, flip angle = 7°, FOV = 256 × 256 mm^2^, and the number of brain slices equal to 176.

### 2.0.2. Data Preprocessing

The resting-state fMRI data are pre-processed using SPM12 (Statistical Parametric Mapping; http://www.fil.ion.ucl.ac.uk/spm/software/spm12/). We discarded first five volumes to allow the magnetization to reach to the steady state value. Next, all functional volumes are slice time corrected using the middle slice as a reference followed by motion correction. Functional scans are spatially normalized onto the Montreal Neurological Institute (MNI) space using the DARTEL procedure, i.e., using the transformation parameters provided by the *T*1-weighted image normalization, resulting in functional images of dimension 53 × 63 × 52 (resampled to 3-mm isotropic voxels). Further, data is smoothed with a Gaussian kernel with 6 mm full width half maximum.

Finally, we regress out nuisance variables (6 head motion parameters, average cerebro-spinal fluid signal from ventricular masks, and average white matter signal from white matter mask) from each voxel’s time series followed by bandpass filtering using a butterworth filter to reduce the low frequency drift and high frequency noise. Hilbert transform is computed for narrowband filtered signals and hence, requires prefiltering of signals. To this end, we bandpass filtered the data in the frequency range of 0.01 to 0.08 Hz. In general, this bandwidth filtering in fMRI is believed to minimize the low frequency drift and the high frequency noise.

Temporal artifacts were identified in each subject’s data by calculating the framewise displacement (FD) from the derivatives of the six rigid-body realignment parameters estimated during motion correction step Power et al. (2014). Subjects with more than 45 brain volumes (30% of total brain volumes) having FD > 0.5 mm were excluded from further analysis (TDC group 4/55; ASD group 12/51). We did not perform scrubbing of data because this may lead to alterations in the temporal structure of data Power et al. (2014).

We included only the male subjects in the study because brain network differences associated with gender may confound differences between the TDC and ASD groups. After quality control, a total of 26 TDC and 35 ASD subjects remained for our analysis. A two-sample t-test with unequal variance showed no significant difference (at *p* < 0.05 significance level) in the age of two groups (Table-1).

**Table 1:** Summary of TDC versus ASD Subjects’ Data

| Characteristic | TDC<br>(S=26) | ASD (S=35) | *p*-<br>value |
| --- | --- | --- | --- |
| Gender | Male | Male |  |
| **Age (years)** |  |  |  |
| Mean (SD) | 10.9 (1.62) | 11.17 (1.49) | 0.50 |
| Range | 8.06–13.79 | 8.25–13.91 |  |
| **Full Scale IQ** |  |  |  |
| Mean (SD) | 120.32<br>(13.53) | 119.06<sup>¶</sup><br>(14.18) | 0.73 |
| Range | 91–148 | 95–149 |  |
| **ADI-R** |  |  |  |
| Social total A | — | 19.74<sup>¶</sup> (5.28) |  |
| Verbal total BV | — | 14.97<sup>¶</sup> (4.91) |  |
| RRB total C | — | 5.09<sup>¶</sup> (2.38) |  |
| R Onset total D | — | 2.56<sup>¶</sup> (1.23) |  |
| **ADOS** |  |  |  |
| Total | — | 10.52<sup>⧧</sup> (4.61) |  |
| Communication | — | 3.18<sup>⧧</sup> (1.54) |  |
| Social | — | 7.33<sup>⧧</sup> (3.53) |  |
| Stereo Behavior | — | 1.89<sup>⧧</sup> (1.58) |  |

TDC: Typically Developing Control; ASD: Autism Spectrum Disorder; ADI-R: Autism Diagnostic Interview-Revised; Social total A: Reciprocal Social Interaction Subscore A; Verbal total BV: Abnormalities in Verbal Communication Subscore; RRB total C: Restricted, Repetitive, and Stereotyped Patterns of Behavior; ADOS: Autism Diagnostic Observation; Stereo Behavior: Stereotyped Behaviors and Restricted Interest. Column on the right displays *p*-values for two sample *t*-test for each sample characteristic. <sup>¶</sup>One subject's score is missing. <sup>⧧</sup>Seven ASD subjects do not have these scores. '—' signifies that these scores are not available for the TDC group.

### 2.0.3. Brain Parcellation

After preprocessing, brain data is parcellated into 90 anatomical predefined ROIs via Automated Anatomical Labeling (AAL) atlas Tzourio-Mazoyer et al. (2002). In this atlas, 45 ROIs lie in the left brain hemisphere and 45 ROIs lie in the right brain hemisphere. Region-representative time series are found for every ROI by averaging time-series of all voxels belonging to the same ROI. This resulted into a matrix X of dimension *T* × *N*, where *T* denotes the number of time points (or the number of times a brain volume is scanned) equal to 147 for a given subject’s fMRI data and *N* denotes the number of ROIs equal to 90 for the AAL atlas.

## 3. Learning and Tracking Dynamic Brain Network Communities

Following steps are proposed for identifying the underlying dynamic brain networks (Fig.1):

Step-1 We utilize phase synchrony based instantaneous measure to compute adjacency matrices of dynamic functional connectivity at each time instant and for all subjects.
Step-2 Tensor decomposition of four-mode tensor (region × region × time × subject) is carried out into components that are used to learn multiple temporal models representing evolution of brain regions over time.
Step-3 *K*-means clustering is used to extract clusters from each learned temporal model.
Step-4 We rank the identified clusters using our proposed cluster scoring metric. Clusters with high scores are labeled as dynamic brain networks that are identified at the group-level.

**Figure 1:**
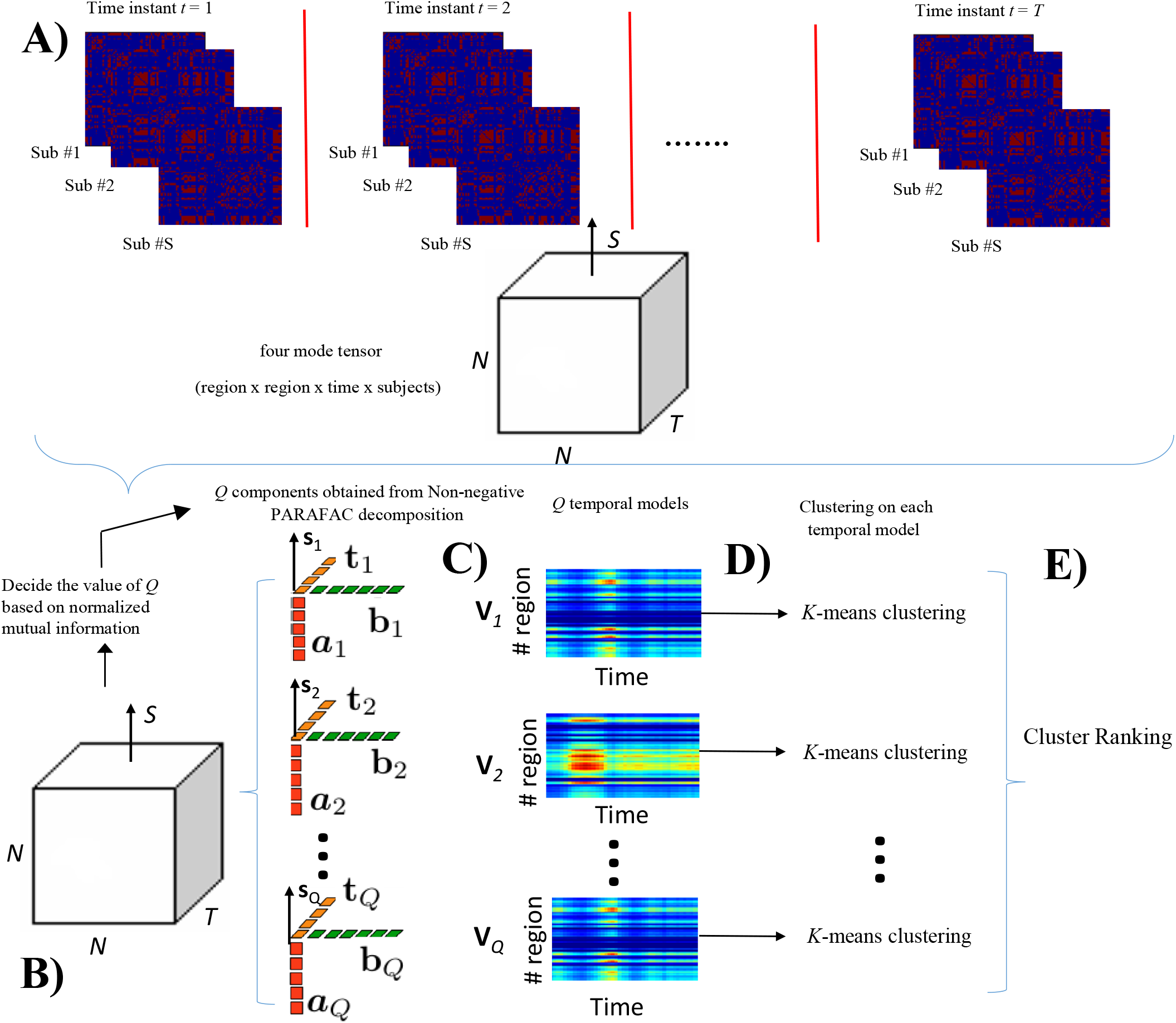
A) High resolution binarized instantaneous coupling matrices are computed for every subject at each time instant, B) Resulting four-mode tensor of dimension *N* × *N* × *T* × *S* is decomposed using non-negative PARAFAC decomposition with *Q* components. C) The temporal model of ROIs in each component is represented by matrix **V**_*q*_ (*q* = 1, 2, …, *Q*). D) *K*-means clustering is used to obtain dynamic brain networks from each temporal model. E) Networks are ranked using our combined cluster score (CCS) presented in Section 3.4 to obtain multiple dynamic brain networks.

### 3.1. Time Varying Phase Synchrony Measure

First, we compute phase synchrony based connectivity matrices of ROIs at all time instants. This allows greater temporal resolution of dynamic functional connectivity compared to the state-of-the-art SW approach.

Hilbert transform of averaged BOLD time series of each ROI is computed to obtain analytic signals, written as *A*(*t*)cos(*φ*(*t*))), where *A*(*t*) and *φ*(*t*) denote the instantaneous amplitude and instantaneous phase, respectively. We removed the first and last ten time points from further analysis in order to reduce border effects caused by Hilbert transform. For any two *i* and *j* ROIs, pairwise phase difference ranged between 0 to τ is computed as:

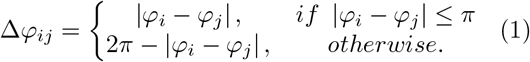

Using the pairwise phase differences between all pairs of ROIs, the instantaneous coupling matrix (ICM) C(*t*), normalized between 0 to 1, is computed as

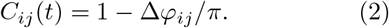

The matrix C(*t*) in the above equation contains values between 0 (no phase matching) and 1 (perfect phase matching). Next, we converted this matrix into sparse, undirected binary matrix using thresholding, as has been done previously in Ponce-Alvarez et al. (2015); Demirta et al. (2016) for phase synchrony based fMRI experiments. Hence, binarized instantaneous coupling matrix C^b^(*t*) from C(*t*) is computed as:

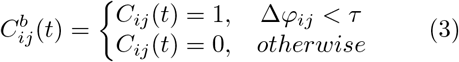

where *τ* denotes a threshold on pairwise phase differences matrix *Δφ_ij_* that is ranged from 0 to *τ*.

#### 3.1.1. Selection of threshold

Recently, in Ponce-Alvarez et al. (2015) positions with pairwise phase difference of more than a threshold value of 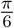 are set to zero in ICM. We have utilized the same value of *τ* in (3) to obtain the adjacency matrix 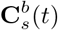 at time *t* for subject *s*. We denote this matrix as the binarized instantaneous coupling matrix (BICM).

A fixed edge threshold has been considered for extracting biologically significant edges in many previous studies (e.g., a threshold of 30% means retaining the highest valued top 30% connections) Achard et al. (2006); Deuker et al. (2009); Zhang et al. (2016). However, on ICM, the 30% edge threshold results in high thresholding value upto ^*π*^ at some time instants. Moreover, it is not necessarily true that a fixed number of edges will be active at all time instants. Hence, in order to avoid any bias from results due to the choice of threshold, all experiments are performed with a fixed threshold of 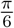 as has been done in Ponce-Alvarez et al. (2015) instead of choosing some fixed number of significant edges. The collection of 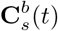 for all subjects *s* = 1, 2, ….., *S* for each group forms a four-mode tensor 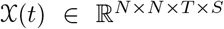 as shown in Fig.1A, where *N* denotes the number of ROIs, *S* denotes the total number of subjects of the corresponding group (TDC or ASD), and *T* denotes the number of time points.

### 3.2. Learning Temporal Models using Subject-Summarized Spatio-Temporal Tensor

We carry out tensor factorization of the 4-mode 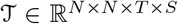 into a linear combination of multiple rank-one tensors to learn multiple temporal models for uncovering the underlying network structures (Fig.1B). This can be achieved by using parallel factor (PARAFAC) decomposition (a.k.a. canonical decomposition) Harshman (1970); Kolda & Bader (2009) as:

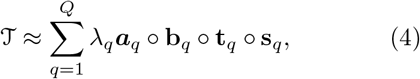

where ‘o’ denotes the outer product and, 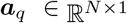 and 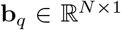 are the loading vectors corresponding to regions *N*, 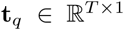 is the time loading vector containing temporal information for tracking the dynamic profile of communities, and 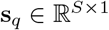 is the subject loading vector. *Q* denotes the number of rank-one tensors (or components) and λ_*q*_ is the singular value corresponding to component *q*. The decomposition in (4) can also be expressed as:

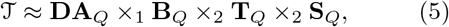

where **D** = *diag*(λ_1_, λ_2_, …λ_*Q*_) and **A**_*Q*_ = [***a***_1_, …, ***a***_*Q*_], **B**_*Q*_ = [**b**_1_, …, **b**_*Q*_], **T**_*Q*_ = [**t**_1_, …, **t**_*Q*_] and **S**_*q*_ = [**s**_1_, …, **s**_*Q*_] are the matrices containing loading vectors of each of the four modes. **A**_*Q*_ (or **B**_*Q*_) in (5) represents ROI specific loading vectors with their corresponding temporal evolution information available in matrix **T**_*Q*_. The column dimension of each of the component matrix is *Q* in the PARAFAC model.

Due to the symmetricity in the adjacency matrix of size *N* × *N* in the first two modes, component matrix **A**_*Q*_ is equal to **B**_*Q*_. Moreover, modes one and two of tensor 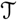 correspond to adjacency matrices with non-negative values. Hence, we add nonnegative constraint on all the four matrices **A**_*Q*_, **B**_*Q*_, **T**_*Q*_, and **S**_*Q*_ in (5). In particular, we carry out non-negative tensor factorization (NNTF) Kim & Park (2012); Cichocki et al. (2009). Implementation of NNTF-based PARAFAC decomposition was based on the non-negative alternate least squares method Paatero & Tapper (1994) combined with a block-coordinate-descent technique Kim & Park (2012), also used recently for fMRI experiments Ponce-Alvarez et al. (2015).

#### 3.2.1. Selection of Q

The value of *Q* in (4) or (5) needs to be specified for PARAFAC decomposition. For each value of *Q*, we extracted communities and compared them with the standard known functional brain networks. We considered a range of 2 to 10 for *Q* and finally, selected *Q* with the highest value of normalized mutual information (NMI).

#### 3.2.2. Computation of Temporal Models

Next, we computed temporal models **V**_*q*_ (*q* = 1, 2, …, *Q*) from every *q*^th^ component using the loading vectors contained in matrices **A**_*Q*_ (or **B**_*Q*_) and **T**_*Q*_, where 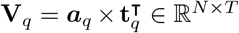. The symbol (.)^**T**^ denotes transpose of a vector and vectors **a**_*q*_ and **t**_*q*_ represent *q*^th^ column of matrices **A**_*Q*_ and **T**_*Q*_, respectively (Fig.1C).

### 3.3. Spatio-Temporal Clustering (STC) Framework

We hypothize that clustering of temporal model **V**_*q*_ along the time dimension (for all *Q* number of components separately) would capture dynamic brain network communities. Thus, first, clusters are estimated from every temporal model **V**_*q*_, *q* = 1, 2, …, *Q* using *K*-means clustering (Fig.1D) Jain & Dubes (1988). We chose *K*-means clustering algorithm for two reasons. First, this problem does not require advanced algorithms beyond *K*-means and second, *K*-means is a fast, easy, and simple clustering algorithm.

A parameter search for the best number of clusters *K* in *K*-means clustering is done using the silhouette criterion index Rousseeuw (1987). The value of *K* that leads to the highest silhouette value is chosen. Silhouette index value is ranged between −1 to 1 with large positive value indicating good clustering. Note that an initial range of *K* is set between 1 to 10 as the maximum number of brain networks, generally available in AAL atlas, is eight such as Visual Network (VN), Auditory Network (AN), Bilateral Limbic Network (BLN), Default Mode Network (DMN), Somato-Motor Network (SMN), Subcortical Network (SN), Memory Network (MN), and Cognitive Control Network (CCN).

We used the Euclidean distance metric for *K*-means clustering and repeated *K*-means algorithm 500 times with random initialization of cluster centers. Clustering result that yielded tightest clusters in terms of Euclidean distance metric was considered for further analysis.

### 3.4. Ranking of Clusters to Obtain Dynamic Brain Network Communities

After obtaining *K_q_* clusters from each of the *q*^th^ component, we require to identify relevant communities from a pool of 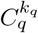 (*q* = 1, …, *Q* and *k_q_* = 1, …, *K_q_*) clusters. We use the term *cluster* to refer to results obtained from STC and the term *communities* to specify ranked and hence, relevant clusters that denote network communities.

To find relevant communities from all available clusters, we create a novel ranking method that is based on two scores: first is the “connectivity strength” and second is the “temporal strength”. Both these scores are combined to define “Combined Cluster Score” in the next subsection that eventually helps with the ranking of clusters.

#### 3.4.1. Connectivity Strength

Connectivity strength of a cluster is related to the strength of connectivity between ROIs belonging to that cluster. A relevant and significant cluster will have a group of tightly interconnected ROIs resulting in a high value of connectivity strength. Owing to this, connectivity strength is considered as an important attribute of cluster score.

First, we compute the connectivity matrix **M***q* using the loading vector 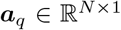 defined in (4) as 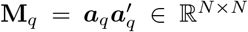. Since we are only interested in ROIs that are part of the given cluster *k_q_*, we compute the desired connectivity matrix 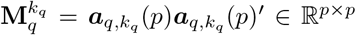, where *p* denotes ROIs that are part of cluster *k_q_* and the corresponding ***a**_*q*_,k_q_* is defined as the *effective loading vector* of regions lying in that cluster.

Each entry 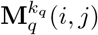 informs about the strength of connection between two ROIs *i* and *j*. We propose to utilize the sum of all entries in this matrix to compute the connectivity strength of cluster *k_q_* and define this as:

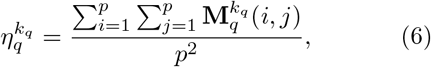

where *q* = 1, …, *Q*, *k_q_* = 1, …, *K_q_*, and division by the total number of entries (a measure of number of regions) is a normalization step with respect to the size of the cluster.

#### 3.4.2. Temporal Strength

Next, we describe the temporal strength score. In general, brain networks are known to be varying over time. Hence, the strength of communities should depend on their temporally varying profile. Thus, the scoring of a cluster should include both the connectivity strength score (6) and the temporal strength score.

To assess the temporal strength of clusters belonging to the *q^th^* component, we utilize the time loading vector **t**_*q*_ of the *q^th^* component to define the temporal strength score of all of its clusters as:

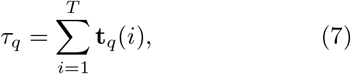

where *q* = 1, …, *Q* and *τ_q_* is the summation of all weight values of *τ_q_* loading vector. In general, a high value of τ_q_ signifies that the region is activated for most of the duration of scan.

#### 3.4.3. Combined Cluster Score

We use the connectivity strength 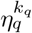 defined in (6) and the temporal strength score *τ_q_* defined in (7) to propose the *Combined Cluster Score* (CCS) of each cluster as:

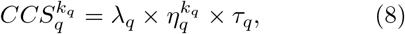

where *q* = 1, …, *Q*, *k_q_* = 1, …, *K_q_*, and *λ_q_* is the PARAFAC decomposition factor of component *q* specified in (4). This is to note that CCS defined above is general and is valid for both the overlapping and non-overlapping communities, although we are detecting overlapping communities.

After computing CCS for each cluster using (8), we generate a ranked list of clusters in the decreasing sorted order of CCS. Clusters at the top of the list are more densely connected compared to the rest of the clusters. Higher ranked clusters provide us the desired dynamic functional brain networks.

## 4. Experiments and Evaluations

In this section, we extract communities that are altered in autism subjects compared to the healthy subjects. First, we computed the phase synchrony based BICM using (3) for the cohort of 26 TDC and 35 ASD subjects obtained from the ABIDE project.

Subject-level ICM and BICM computed at four random time points of one randomly chosen subject from both TDC and ASD groups are shown in Fig.2. It is difficult to find differences between the two groups by visualizing these matrices, representing regional modular connectivity.

**Figure 2:**
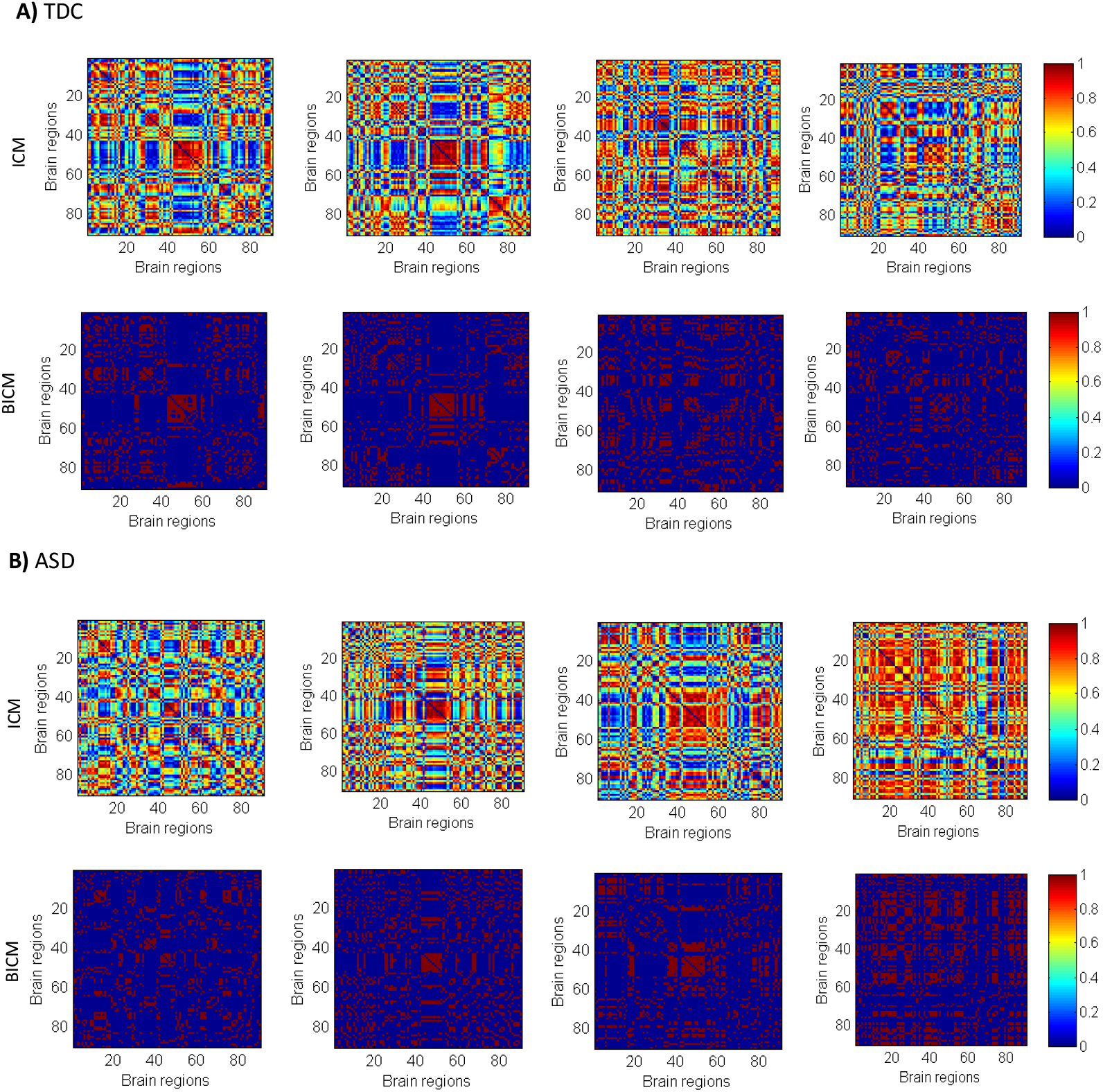
The ICM and BICM at four (randomly chosen) time points (10, 45, 90, and 120) on one subject chosen randomly from both the groups: A) TDC B) ASD.

### 4.1. High Resolution Temporal Models

We applied NNTF-based PARAFAC decomposition on four-mode tensor 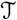 of both the groups (TDC and ASD) and obtained *Q* temporal models, as discussed in section 3.2. We chose an initial value of *Q* in the range of 2 to 10. We observed the highest NMI value at *Q*=10 for both the groups and hence, factorized tensor 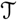 into *Q*=10 components.

A typical randomly chosen regional loading vector ***a**_*q*_* is shown in Fig.3A and its temporal profile ***t**_*q*_* is displayed in Fig.3B for both the TDC and ASD groups. The bottom panel of Fig.3 shows the corresponding temporal model **V***_q_* depicting interregional connectivity strength changing over time.

**Figure 3:**
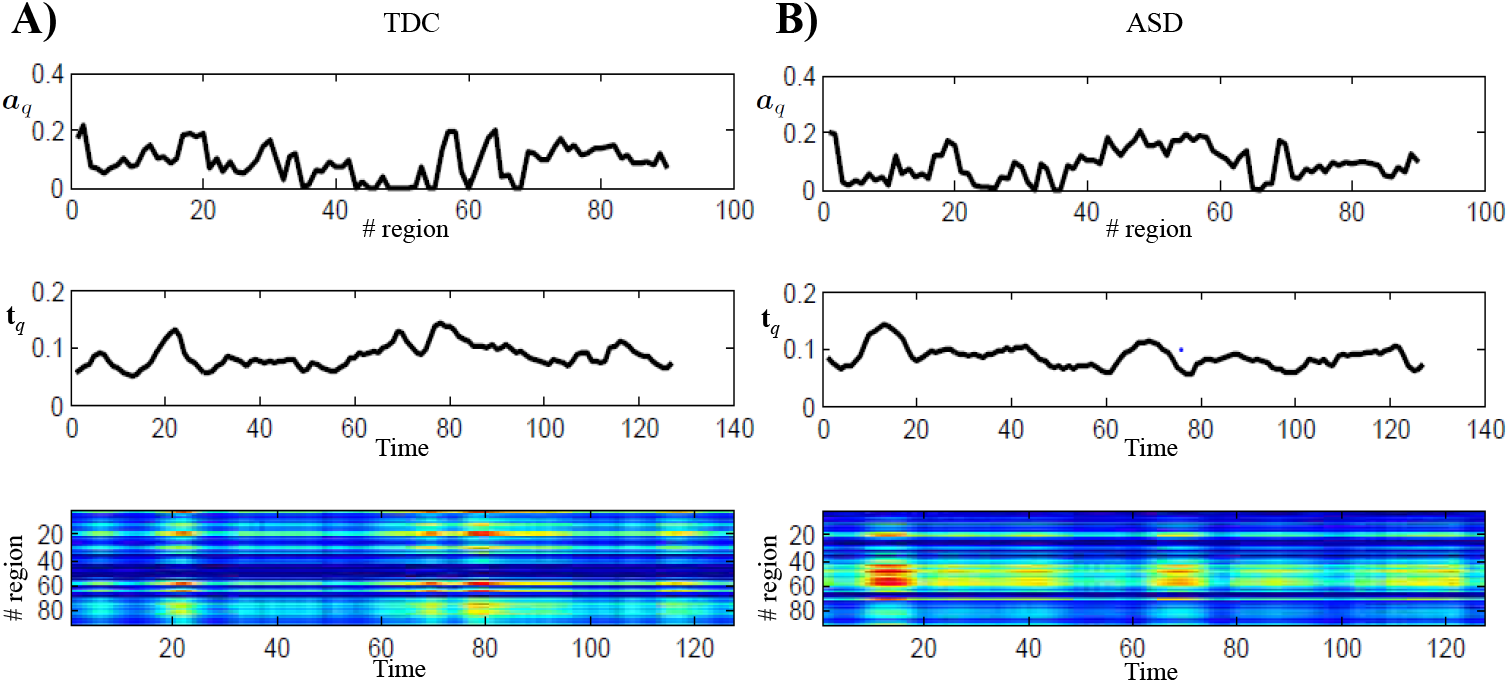
Plot of one randomly chosen regional loading vector ***a**_q_*, its temporal loading vector ***t**_q_*, and the corresponding matrix **V***_q_* for both A) TDC and B) ASD groups.

### 4.2. Extraction of Statistically Different Components between ASD and TDC Groups

In order to identify altered components of ASD compared to TDC, we define the strength of each component using the temporal, subject and region’s loading vectors as 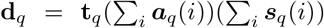. The formulation of strength of components for three-mode tensor was first proposed in social networks Gauvin et al. (2014). We extended this for the case of 4-mode tensor, where fourth mode of tensor contains information about subjects. We used two-sample *t*-test (at *p* < 0.05) on the strengths of components of TDC and ASD groups for identifying components that are statistically different between the two groups. We also corrected for multiple comparison using false discovery rate (FDR) at α= 0.05 Benjamini & Hochberg (1995). This process is shown in Fig. 4 and the results of this statistical test are shown in Table-2.

**Figure 4:**
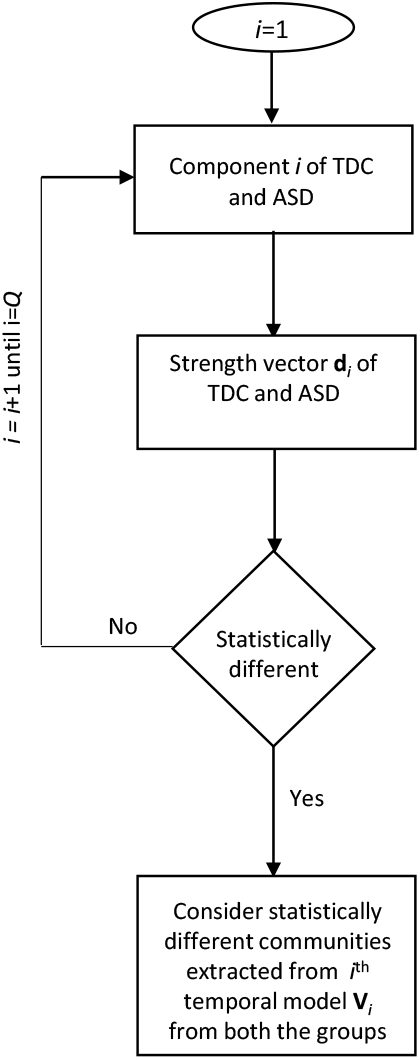
Identification of statistically different components between TDC and ASD groups using the strength vectors of components.

**Table 2:** Identified phase synchrony based statistically different components (marked with bold *p*-values) between the two groups (two-sample *t*-test, FDR corrected *p* < 0.05)

|  | Component# | 1 | 2 | 3 | 4 | 5 | 6 | 7 | 8 | 9 | 10 |
| --- | --- | --- | --- | --- | --- | --- | --- | --- | --- | --- | --- |
| TDC | Number of Communities | 8 | 2 | 2 | 2 | 6 | 2 | 2 | 2 | 2 | 2 |
| ASD | Number of Communities | 10 | 2 | 2 | 2 | 2 | 2 | 2 | 2 | 2 | 8 |
| | $p$-value of $t$-test on strength vectors of corresponding components of ASD and TDC groups | **2.51e-21** | **1.48e-26** | 0.08 | **2.48e-21** | **8.72e-25** | 0.01 | **1.14e-05** | 0.024 | **0.06e-02** | 0.04 |

### 4.3. Community Structure and the Associated Temporal Activity Patterns

We considered only those temporal models **V**_*q*_ that corresponded to statistically different components across ASD and TDC groups. We applied *K*-means clustering on each of these selected temporal models separately to identify multiple clusters corresponding to each temporal model. Optimum value of *K* is decided based on the maximum value of Silhouette index. From Table-2, we note that components#1, 2, 4, 5, 7, and 9 are statistically different (FDR corrected, *p* < 0.05) between the two groups. However, the number of communities between these components except for component#1 are few. Since the number of functional brain networks are around 7 to 10, it shows that these components are doing random grouping of regions in a few communities. Hence, only component#1 is considered to be different between the two groups.

Clusters/communities of component#1 are ranked with the proposed CCS score in (8), where different ranking of the same region between groups (TDC versus ASD) and clustering with different regions between groups specify differences in activation and connectivity patterns between the two groups, respectively.

Tables-3 and 4 illustrate the region-level description of communities obtained from component#1 of both the groups. We list communities according to the decreasing order of the CCS score. Fig.5 presents these communities on the axial brain slice. Here, we also present ground-truth labels of communities present in the AAL atas for comparison. The temporal evolution of these communities in both the groups is presented in Fig.6.

**Figure 5:**
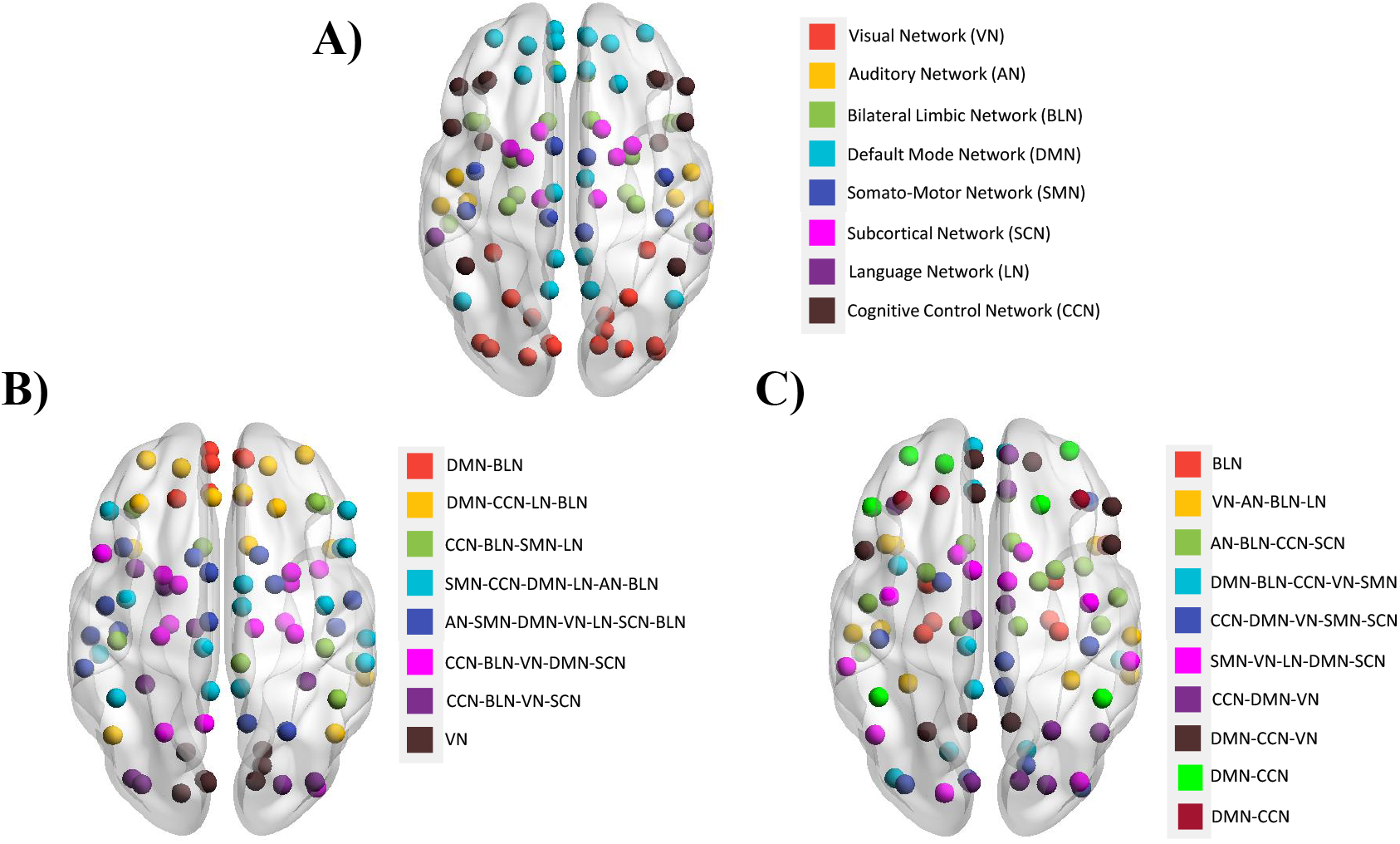
List of communities in A) AAL atlas, B) TDC group (component 1-8 communities), and C) ASD group (component 1-10 communities). We have presented TDC and ASD communities according to their CCS ranking score.

**Figure 6:**
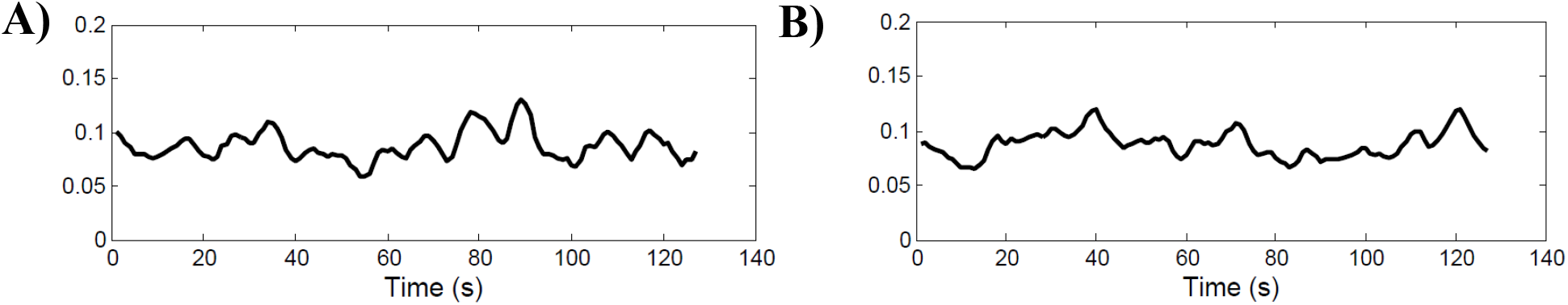
Temporal dynamics of component#1 in both the groups: A) TDC, B) ASD.

Our results on TDC group in Table-3 reveal that the highest CCS community is comprised of 4 ROIs (community 1 in Table-3) containing the left dorsal, medial, medial orbital superior frontal gyrus, and gyrus rectus, forming a network associated with DMN and BLN. In general, these regions are strongly active in healthy brain during the rest condition Monk et al. (2009). On the other hand, ASD group results in 4 bilateral limbic regions (hippocampus, parahippocampal gyrus, amygdala and right temporal pole superior temporal gyrus) that are clustered in the first community. Previous studies also reported hyperconnectiviy in the right parahippocampal gyrus of ASD compared to the TDC group Monk et al. (2009).

**Table 3:**
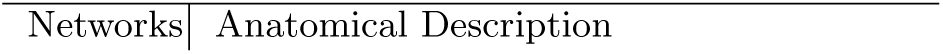
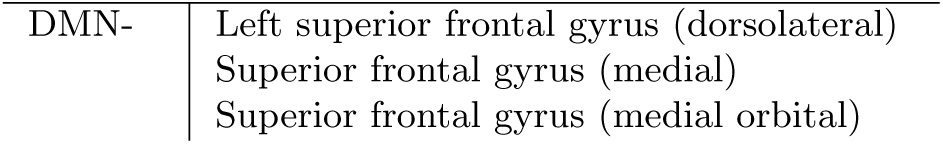
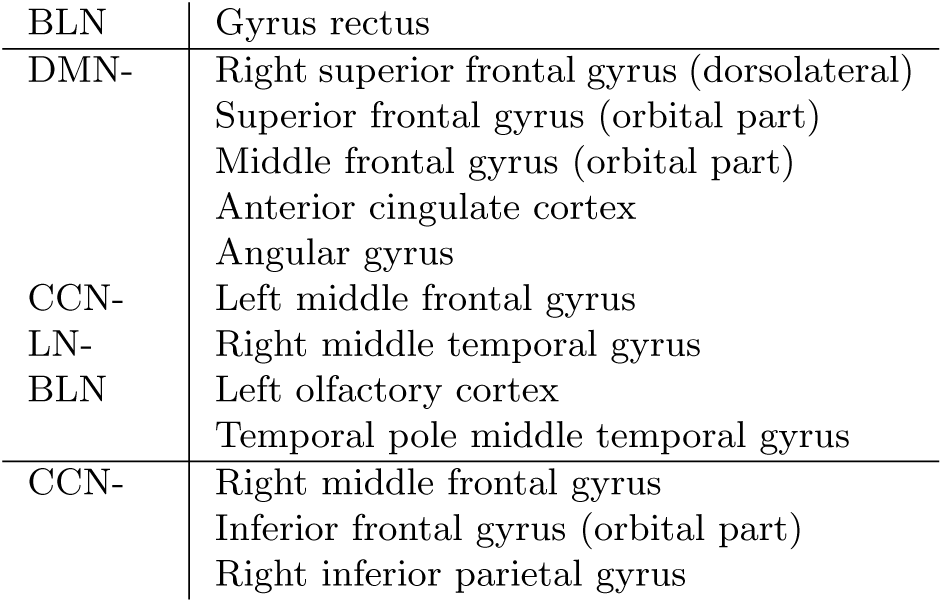
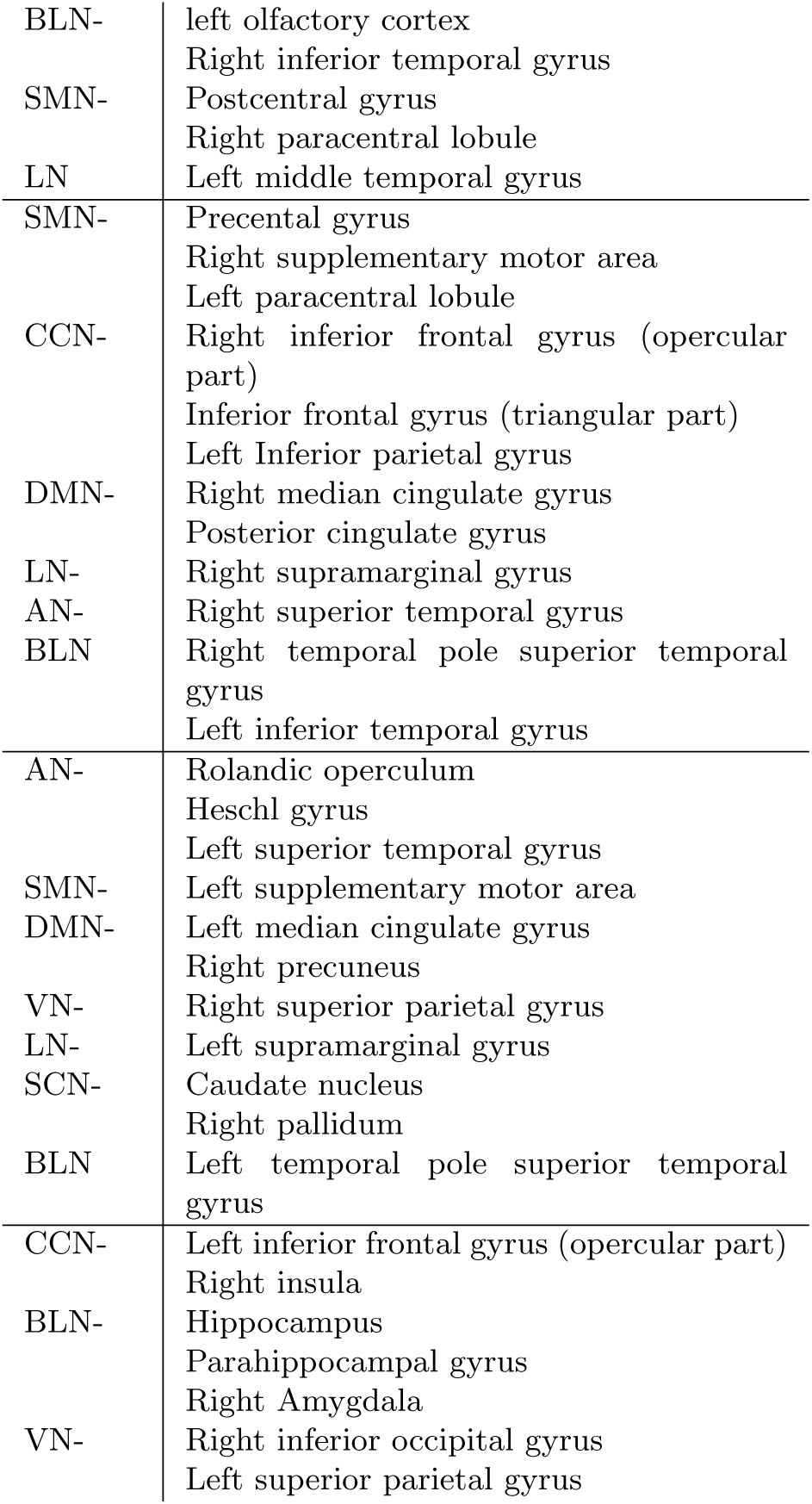
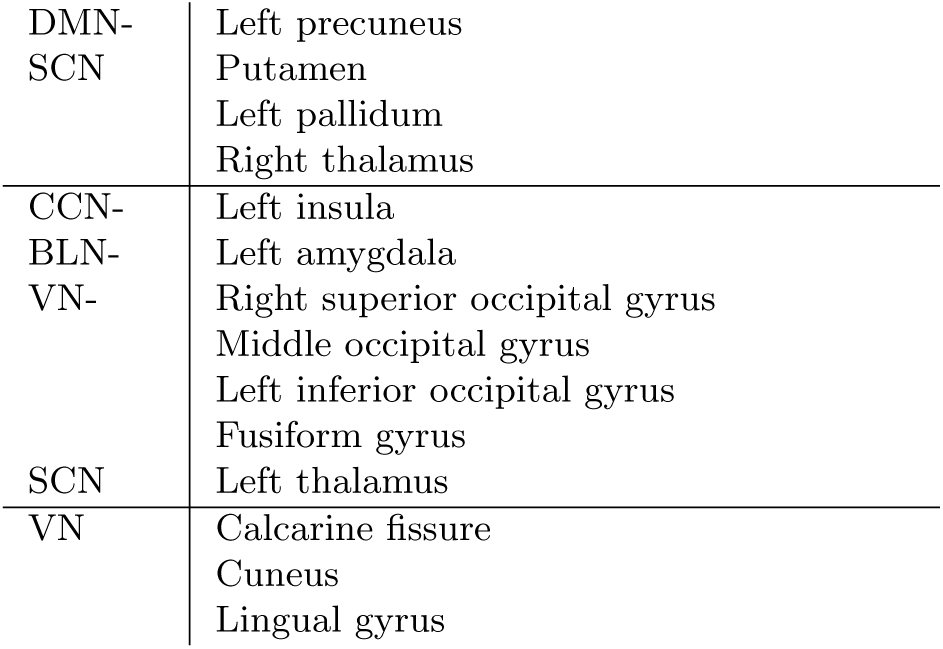
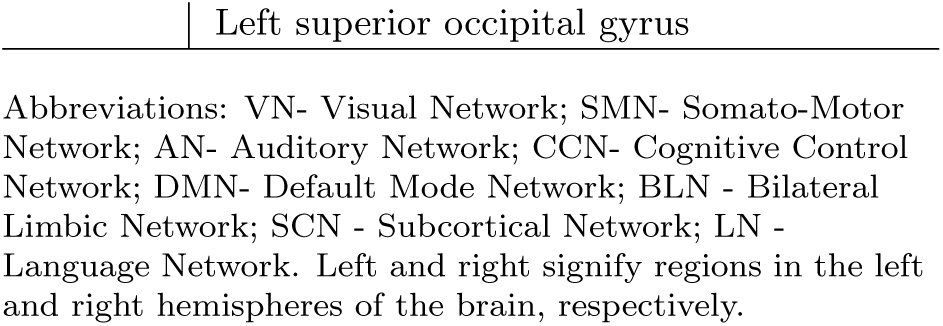
List of regions forming communities in the TDC group for component # 1

**Table 4:**
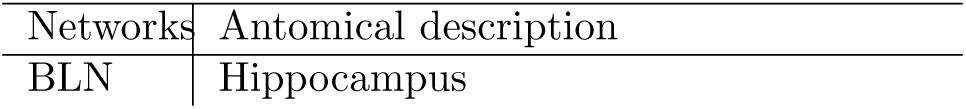
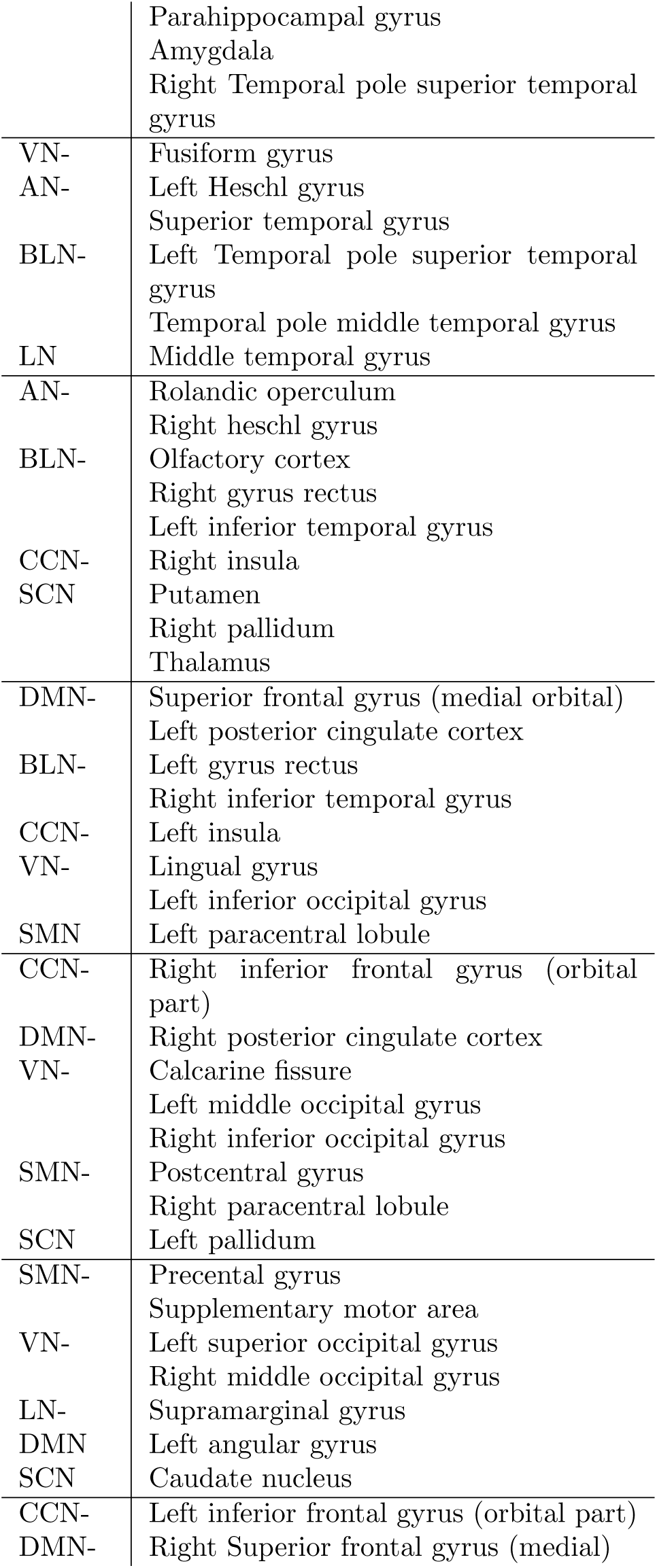
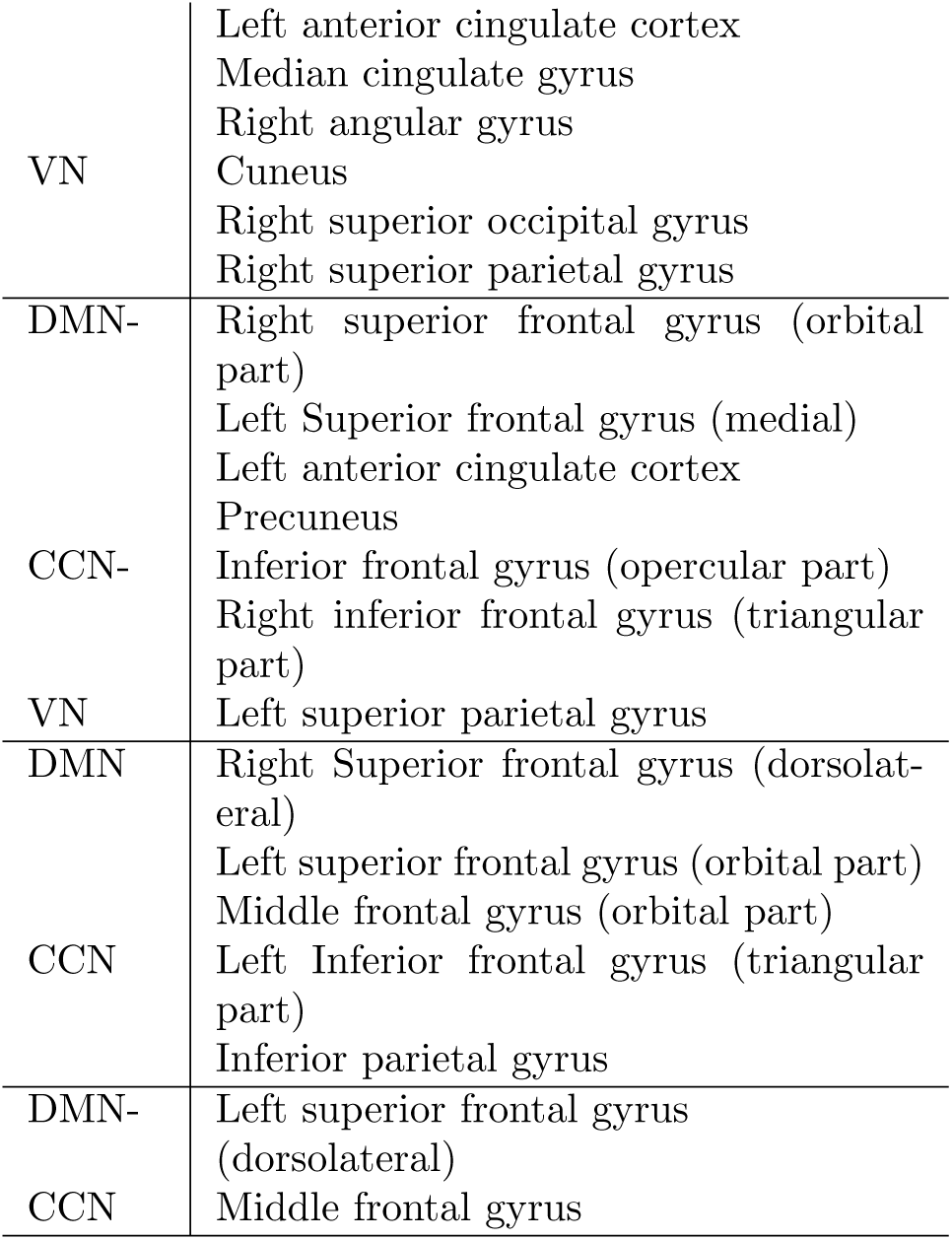
List of regions forming communities in the ASD group for component # 1

**Table 5:** Lowest ranked altered brain regions in ASD compared to TDC

| Networks | AAL regions |
| --- | --- |
| VN | Superior parietal gyrus |
| BLN | Left olfactory cortex<br>Gyrus rectus<br>Right inferior temporal gyrus |
| DMN | Superior frontal gyrus<br>Middle frontal gyrus (orbital part)<br>Anterior cingulate cortex<br>Median cingulate gyrus<br>Posterior cingulate cortex<br>Angular gyrus<br>Precuneus |
| SMN | Precentral gyrus<br>Supplementary motor area<br>Posterior gyrus<br>Right paracentral lobule |
| SCN | Caudate nucleus |
| LN | Supramarginal gyrus |
| CCN | Middle frontal gyrus<br>Inferior frontal gyrus<br>Inferior parietal gyrus |

Next, we present the lowest ranked regions of all networks in ASD compared to TDC group in Table 5. From this table, we observe that the proposed method could find 20 altered lowest ranked regions in ASD group (two sample *t*-test, FDR corrected *p* < 0.05). These regions belong to VN (superior parietal gyrus), BLN (left olfactory cortex, gyrus rectus, right inferior temporal gyrus), DMN (superior frontal gyrus, orbital part middle frontal gyrus, anterior and posterior cingulate cortex, median cingulate cortex, angular gyrus, and precuneus), SMN (precentral and posterior gyrus, supplementary motor area, right paracentral lobule), SCN (caudate nucleus), LN (supramarginal gyrus), and CCN (middle frontal gyrus, inferior frontal gyrus, and inferior parietal gyrus).

## 5. Discussion

In this work, we have attempted to develop a better understanding of dynamic fluctuations occurring in the resting-state FC of diseased and normal subjects. FC is often computed and interpreted within some fixed time duration. These are known as static brain networks. Recently, results have been reported on the instantaneous interpretation of dynamic FC computed by utilizing the phase difference between pairs of voxels’ time series. This phase difference is one of the ways to measure temporal dynamics between regions and is commonly known as phase synchrony.

We examined dynamic reconfiguration of functional brain networks via our proposed method based on ICM, generated from phase synchrony. These matrices are thresholded to preserve the small, but significantly relevant phase synchrony range. Next, the 4-mode tensor is decomposed using NNTF to build multiple temporal models. This is followed by *K*-means clustering and the ranking of resulting clusters via the proposed CCS measure. This complete process yields appropriate communities. We built communities for both the healthy and autistic subjects to understand the probable alterations of networks in neuro-disorders.

### 5.1. Community Structure of dFC States in Healthy Brain

By learning and tracking dynamic brain network communities (Section 2), we find many clusters of commonly reported functional brain networks including the default mode network, visual, cognitive, somato-motor, auditory, bilateral limbic, subcortical, and language networks (Table-3). This suggests that binarized instantaneous coupling matrices of phase synchrony contain meaningful information about the large scale brain networks of healthy individuals within 0.01 to 0.08 Hz. Hence, we propose a novel framework of modeling and interpreting dFC that are evolving over time.

We observe that the resting-state communities on TDC, transiently emerge and dissolve in time (Fig. 6), giving rise to dynamic reconfiguration of brain networks. Consistent with our findings, temporal fluctuations of dFCs have also been reported earlier using sliding window analysis of fMRI signals on healthy subjects.

### 5.2. Within Group Differences in Autism

ASD causes abnormality in functional brain networks via a variety of underlying genetic and acquired causes Yao et al. (2016). Thus, study of brain networks in ASD is a key for understanding the altered functioning in the diseased brain. In order to meet this goal and to have a better understanding of altered brain networks in ASD, we learned the community structures of communities in the ASD group (Section 2).

Our analysis suggests that compared to the TDC group, autism subjects show reduced ranked (and hence, weakly connected) communities of VN, BLN, DMN, SMN, SCN, LN, and CCN.

### 5.3. Characterizing Regions Distinctive in Autistic Disorder

We carried out detailed investigation on the communities in both the groups. In particular, we want to know regions observed with less ranking in ASD, i.e., hypo-active regions in ASD.

#### 5.3.1. Hypo-active Regions in ASD

Hypo-active regions include the VN (superior parietal gyrus), BLN (left olfactory cortex, gyrus rectus, right inferior temporal gyrus), DMN (superior frontal gyrus, orbital part middle frontal gyrus, anterior and posterior cingulate cortex, median cingulate cortex, angular gyrus, and precuneus), SMN (precentral and posterior gyrus, supplementary motor area, right paracentral lobule), SCN (caudate nucleus), LN (supramarginal gyrus), and CCN (middle frontal gyrus, inferior frontal gyrus, and inferior parietal gyrus).

### 5.4. Limitations and Future Directions

The method developed in this work allows us to extract a group level using undirected phase synchrony measure. However, directionality aspect of brain networks should also be exploited for complete understanding of diseased brain networks. Directed adjacency matrix weights can be estimated using time-shifted version of region time series, as opposed to dFC.

Correlation of Hilbert envelopes also provides information about functional connectivity O'Neill et al. (2017). However, in this work, we have discarded the magnitude of the analytic signals, also known as Hilbert envelope. The examination of Hilbert envelope based correlations might bring new important findings about brain networks and hence, in future, we would like to utilize both envelope and phase information of Hilbert transform for building more robust brain networks.

In Autism, although brain regions show activation on internal or external stimuli, there is altered connectivity between regions and hence, the functional brain networks are affected. This problem leads to reduced attention, difficulty in prioritizing tasks, inhibition in brain required for executing intended tasks, and behavioral problems as stated in the previous section. These problems add difficulty for their caretakers as well. The work carried out in this paper on dynamic functional brain networks can be utilized to develop interventions for these subjects as part of neuro-rehabilitation. We plan to take up this challenge as the future work.

## 6. Conclusions

In conclusion, a novel methodology has been proposed for estimating dynamic brain network communities that are computed at every time instant. We present application of the proposed framework on ASD subjects demonstrating altered dynamic brain network communities in autism subjects compared to the TDC group. This further supports the hypothesis that autism is a disorder affecting brain networks. We observe that the proposed methodology is able to report changes in large scale networks such as DMN, VN, BLN, SMN, SCN, LN and CCN. Impaired connectivity in the resting state brain networks is due to the impairment in neuropsychological function that suggests that study of communities is a potential biomarker in ASD. We have observed reduced ranked activation in different communities. This suggests that instantaneous functional connectivity can play a key role in the study of reorganization of brain networks.

## Acknowledgements

The first author would like to thank Visvesvaraya research fellowship, Department of Electronics and Information Tech., Ministry of Comm. and IT, Govt. of India, for providing the financial support for this work. We also thank ABIDE for sharing the data online.

